# DeepCapTail: A Deep Learning Framework to Predict Capsid and Tail Proteins of Phage Genomes

**DOI:** 10.1101/477885

**Authors:** Dhoha Abid, Liqing Zhang

## Abstract

The capsid and tail proteins are considered the main structural proteins for phages and also their footprint since they exist only in phage genomes. These proteins are known to lack sequence conservation, making them extremely diverse and thus posing a major challenge to identify and annotate them in genomic sequences. In this study, we aim to overcome this challenge and predict these proteins by using deep neural networks with composition-based features. We develop two models trained with *k*-mer features to predict capsid and tail proteins respectively. Evaluating the models on two different testing sets shows that they outperform state-of-the-art methods with improved F-1 scores.

## Introduction

Phages or bacteriophages are viruses that infect bacteria. These microorganisms can reproduce through two different life cycles, lysogeny and lytic. For the lysogeny cycle, the phage integrates its genome with the bacteria genome and stays there. In this cycle, the phage becomes part of the bacterial genome and replicates together with the bacteria; whereas for the lytic cycle, the phage enters the bacterial cell, uses its machinery to replicate, reproduce new phages, and then lyses the cell membrane to disperse into the environment, resulting in death of the invaded bacterium [1].

Phages are getting increasing attention primarily due to the advent of the shotgun metagenomic sequencing. This technology enables comprehensive sampling of the genomes that are present in a given environmental sample, such as soil and seawater, while circumventing the culture of the microorganisms, which is both labor intensive and often infeasible [2]. Furthermore, there is increasing interest in characterizing the interactions between phages and their bacterial hosts. Understanding the interactions has important implications, one of which is in combating antibiotic resistance in bacteria where phages can be introduced to infect and kill pathogenic bacteria or induced into lytic stage if already integrated in bacteria [3].

However, uncovering viral sequences has been challenging. Despite being the most abundant organisms on earth with an estimate of more than 10^30^ [4], only 8108 complete virus genomes are curated at NCBI currently. Consequently, methods for predicting/annotating viral sequences that rely heavily on reference databases, such as the alignment-based methods, are not effective in detecting novel viruses and phages. Indeed, if the input sequence does not align to any sequence in the reference database, it would be annotated as unknown sequences. This problem is further exacerbated by the lack of well established taxonomic and phylogenetic relationships in viruses and phages as they do not have the ribosomal genes that are conserved universal markers in other organisms for phylogenetic classifications [5].

To overcome these challenges, composition-based methods were introduced to circumvent the requirement of universal or markers genes [6–8]. The composition-based methods use the composition of the sequence, such as *k*-mers, as features to train machine learning models and then use the trained models to predict the taxonomy of new sequences. The composition-based prediction methods can make prediction on any sequence even if it does not align to the reference database.

Many studies built machine learning models to classify a given sequence to either viral or nonviral sequences. For example, Feng et al. trained a naive Bayes classifier with amino acid composition and dipeptide composition as features [9]. Later, Ding et al. developed PVPred that uses SVM with g-gap dipeptide composition as features and selects the most significant features by analyzing the variance [10]. Subsequently, Zhang et al. developed a random forest ensemble method with a set of features that include pseudo-amino acid composition (PseAAC) and position-specific scoring matrix (PSSM) [11]. Recently, Manavalan et al. used SVM with feature selection of amino acid composition to classify sequences as viral or nonviral proteins [6].

Other studies focus on models that classify phage protein sequences to either structural or nonstructural proteins. Structural proteins can be either capsid or tail proteins, and nonstructural proteins can be anything else such as viral binding or even bacterial proteins. Capsid and tail proteins are considered the footprint of the phage genome. The tail protein encodes a tail shape protein that acts as a mediator for the phage to attach onto the membrane of the bacteria during infection. The capsid protein encodes a shell structure that encapsulates the phage genetic material. The capsid is known to exist in all sequenced viruses and phages; it protects the phage/virus from degradation by the host enzymes. The capsid also acts as a mediator to infect bacteria by attaching the phage to its host and enabling its penetration through the host membrane.

These vital roles of capsid and tail proteins motivated Seguritan et al. [7] to develop iVIREONS to predict them. iVIREONS consists of a set of 30 artificial neural network models that use amino acid frequency and isoelectric as features. However, the set of models can output different predictions for the same input, which challenges the user to determine the correct prediction. More recently, another machine learning model VIRALpro was developed to also predict capsid and tail sequences [8] using SVM with amino acid frequency and HMM models. VIRALpro outperforms iVIREONS [8], but is remarkably slower since it uses HMM.

In this study, we built two machine learning models that also predict capsid and tail proteins respectively. For this purpose, we used the deep neural network models; these models are considered the most modern machine learning models to date, known for their exceptional performance that outperform their predecessors [12]. They have been extensively used in the fields of computer vision and natural language processing [12], and only recently gained attention in the field of genomics [13]. However, to our knowledge, there has not been any study that harnesses the power of these models to predict capsid and tail proteins.

We propose two distinct deep neural network models that predict capsid and tail phage protein respectively. We trained the models using *k*-mer frequency as features and examined different *k*-mer sizes ranging from one to four. We evaluated the models with two test data sets and compared our models with iVIREONS [7] and VIRALpro [8].

## Materials and Methods

### Data Collection

We collected all the phage and prophage sequences from Phaster [14]. The Phaster database consists of curated phage and prophage proteins taken from NCBI and the prophage database [15]. This database is publicly available and regularly updated.

As of May 2018, Phaster included a total of 260, 403 phage and prophage protein sequences. Redundant sequences (i.e., sequences are identical to each other) were removed, leaving 187, 670 unique sequences. Figure 1 shows the distribution of the protein sequences of Phaster after removing the redundant sequences: 66.38% are hypothetical or putative proteins, 9% are enzymes, and 16.31% are hard to categorize with no clear description. The remaining 8% consist of capsid, tail, and nonstructural proteins, and are used in this study.

**Fig 1.**
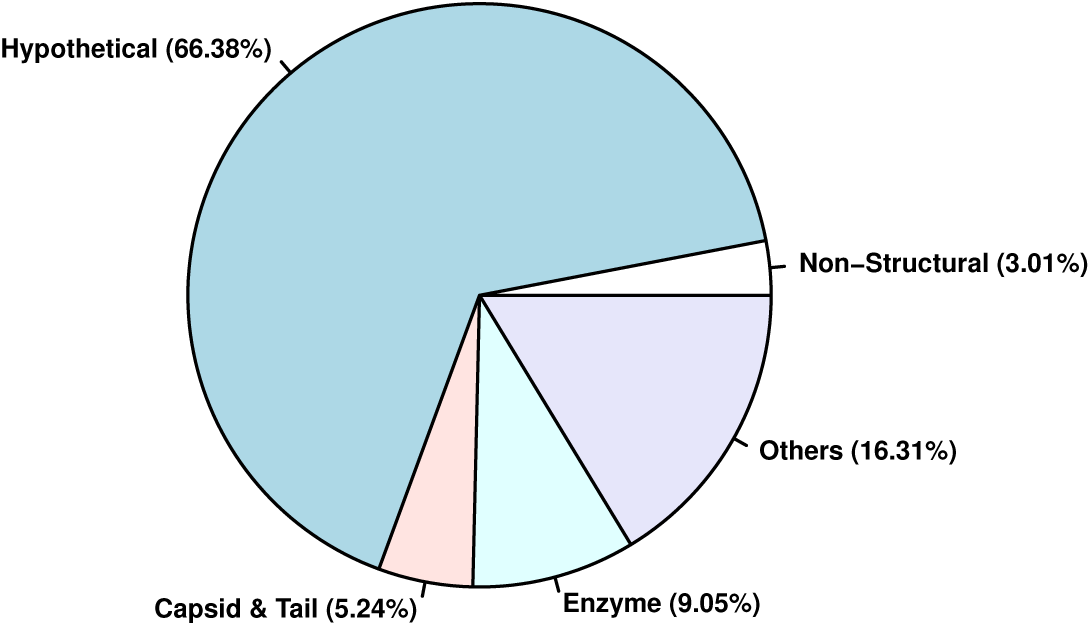
Distribution of protein sequences in Phaster database.

The capsid, tail, and nonstructural proteins were annotated similarly to iVIREONS [7] and VIRALpro [8], that is, the description of the fasta files was used to annotate the proteins. For example, if the description contains the word ‘capsid’, the corresponding sequence was labeled as capsid protein. Hypothetical or putative sequences were not included to ensure data quality. A comprehensive list of the words used for the annotation is provided in Table 1 of the supplemental material. Thus the annotated proteins include 3, 401 unique capsid proteins, 6, 442 unique tail proteins, and 5, 654 unique nonstructural proteins. These proteins are divided into training and testing sets, as detailed in the next section.

**Table 1.** Distribution of capsid, tail, and nonstructural protein sequences in the training and testing sets for capsid and tail models.

| Dataset | # of sequences to train & test for capsid models | # of sequences to train & test tail models |
| --- | --- | --- |
| **Training set** | Capsid (P): 2241<br>Nonstructural (N): 3678 | Tail (P): 4229<br>Nonstructural (N): 3678 |
| **Representative testing set** | Capsid (P): 960<br>Nonstructural (N): 1576 | Tail (P): 1813<br>Nonstructural (N): 1576 |
| **Independent testing set** | Capsid (P): 200<br>Nonstructural (N): 400 | Tail (P): 400<br>Nonstructural (N): 400 |

(P) means positive sequences, (N) means negative sequences. The same nonstructural sequences are used as negative sequences for both capsid and tail models.

### Training & Testing sets

The capsid, tail, and nonstructural proteins were split into three sets, one training set and two testing set. The three sets are mutually exclusive, the same protein sequence does not exist in more than one set. The training set is used to train the deep learning models and the two testing sets are used to validate the trained models. We call the first testing set the *representative* testing set because it includes sequences that are similar to the training set. Figures 2a and 2b show the identity of the best-hits of the representative testing set against the training set. These figures show that the majority of sequences in the representative testing set are highly similar to the training set: 80% of these sequences have an identity between 40% and 100%. The second testing set is the *independent* testing set, including proteins that are less similar to the training set. Figures 2c and 2d show the identity of the best-hits of the independent testing set against the training set: 60% of the sequences in the independent testing set have an identity of less than 20%, and more than 30% have an identity between 20% and 40%. These best-hits are generated by blastp: the testing set, whether the representative or the independent set, is the query, and the training set is the subject.

**Fig 2.**
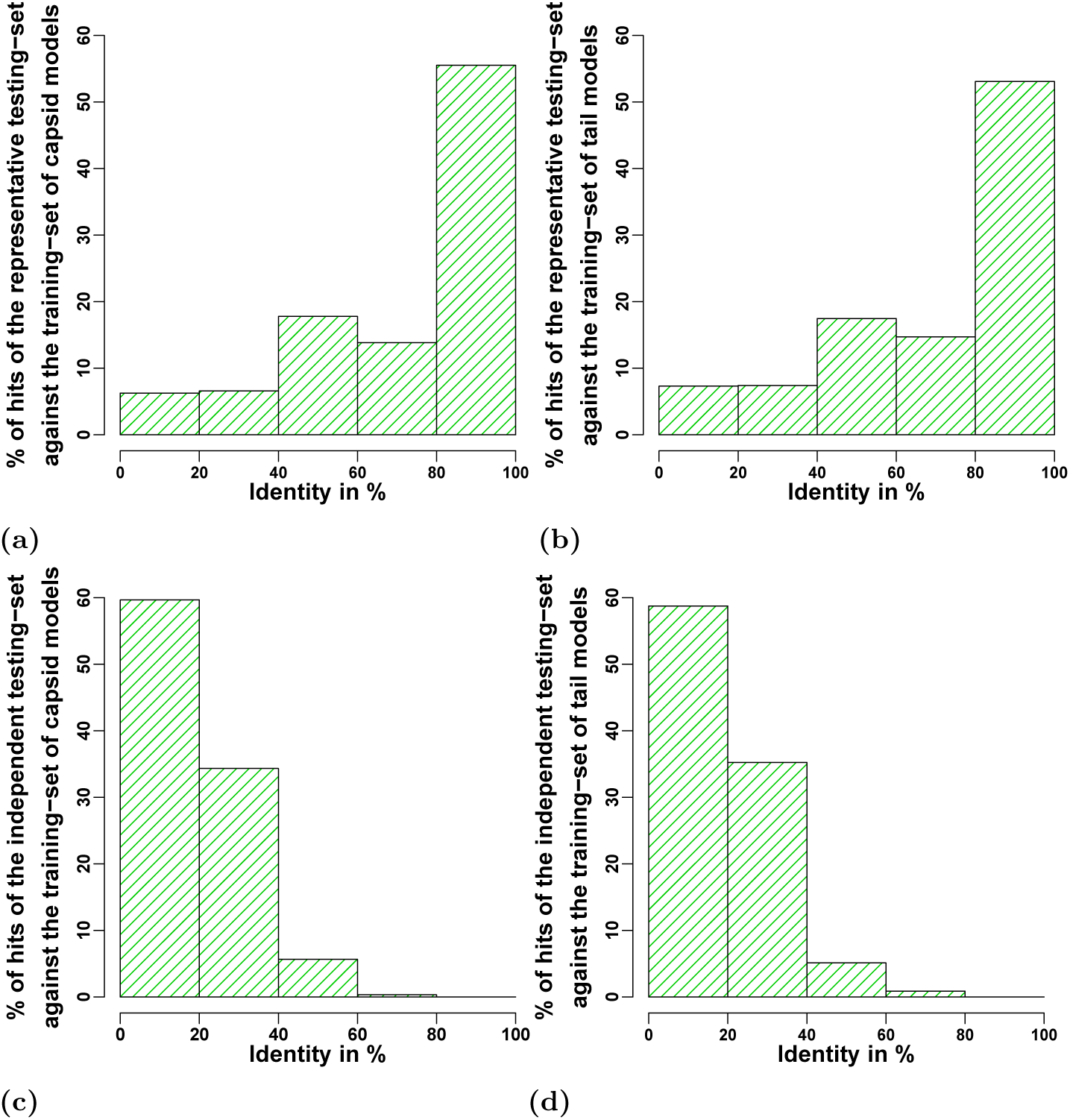
Distribution of identity of the best-hits of the testing and training sets. Figures 2a and 2b present the best-hits of the representative set against the training set for capsid and tail models respectively, and Figure 2c and 2d show the best-hits of the independent set against the training set for capsid and tail models respectively.

Table 1 shows the number of sequences used in the training and testing sets for capsid and tail models. The training and testing sets are selected randomly (see supplemental Figure 1 for details on how we built these sets).

### Extraction of *k*-mer Features

*K*-mer frequencies of the protein sequences were used as features to train the different deep learning models. Various *k*-mer sizes were examined ranging from one to four. Table 2 shows the number of features for every *k*-mer size (e.g., for *k*-mer size ≤ 2, we would have 420 features with 20 features for *k*-mer size one and 400 features for *k*-mer size two).

**Table 2.**
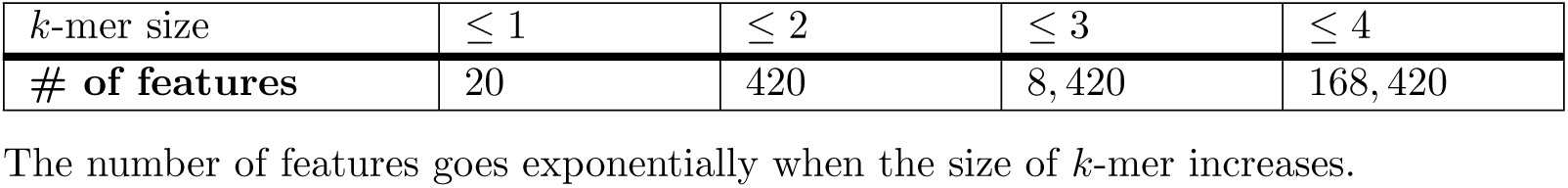
number of features based on the *k*-mer size.

| $k$-mer size | $\leq 1$ | $\leq 2$ | $\leq 3$ | $\leq 4$ |
| --- | --- | --- | --- | --- |
| # of features | 20 | 420 | 8,420 | 168,420 |

The number of features goes exponentially when the size of $k$ -mer increases.

### Training Deep Neural Network Models

Four different deep learning architectures with varying numbers of layers and nodes were investigated: the first model has two layers with 12 and 8 nodes respectively; the second model has four layers with 400, 200, 100, and 50 nodes respectively; the third model has four layers with 600, 300, 150, and 60 nodes respectively; the fourth and most complex model has four layers with 800, 400, 200, and 100 nodes respectively.

For all models, the following parameters were used: “relu” as activation function, “adam” as optimizer, and “binary crossentropy” as loss function. The models were trained using 150 epochs with a batch size of 10. The architectures as well as the parameters used were determined through extensive experimentation.

We provide a naming convention for these models in Figure 3. For example, ‘*Capsid:12:8*’ predicts capsid proteins and has two layers: 12 nodes in the first layer and eight nodes in the second layer. ‘*Tail:400:200:100:50*’ predicts tail proteins and has four layers: 400 nodes in the first layer, 200 nodes in the second layer, 100 nodes in the third layer, and 50 nodes in the forth layer.

**Fig 3.**
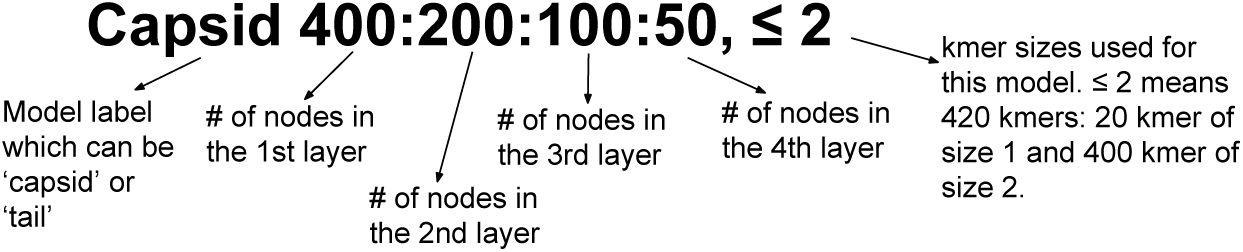
Naming convention of the implemented deep neural network models.

### Validation of The Trained Models

#### Two Different Testing sets

To validate the trained models, two different testing sets were used: the representative and independent testing sets. The representative testing set assesses the models when the input sequence is similar to the training set. On the other hand, the independent testing set evaluates the models when the predicted sequence is considerably different from the training set, which mimics the real-world problem of virus annotation where a significant number of new sequences are not similar to the reference database sequences.

#### Performance Criteria

Accuracy, F1-score, Recall, and Precision were used to assess the prediction of the trained models. We present the formula of Accuracy, F1-score, Recall, and Precision in Equations 1, 2, 3, and 4 respectively:

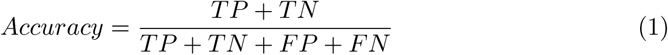

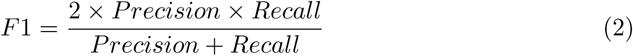

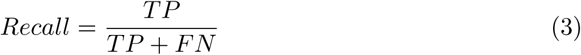

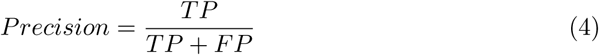

TP is the number of capsid or tail sequences that are classified correctly, TN is the number of the nonstructural sequences that are classified correctly, FP is the number of the nonstructural sequences that are classified incorrectly (as either capsid or tail), FN is the number of capsid or tail sequences that are classified incorrectly (as nonstructural).

## Results

### Framework of The Proposed Predictor

DeepCapTail is a publicly available framework that predicts capsid and tail proteins. This framework can be downloaded at https://github.com/Dhooha/DeepCapTail. This framework consists of a machine learning project written in Python and uses the scikit-learn library [16]. Figure 4 shows the steps followed to build DeepCapTail: (1) different deep neural network architectures were investigated with different *k*-mer sizes in order to decide on the most effective architectures as well as *k*-mer sizes; (2) the most effective deep neural networks were used to train capsid and tail models using the training set; (3) these models were tested using two distinct testing sets, dubbed representative and independent testing sets; (4) the best capsid and tail models were selected based on the F1-score.

**Fig 4.**
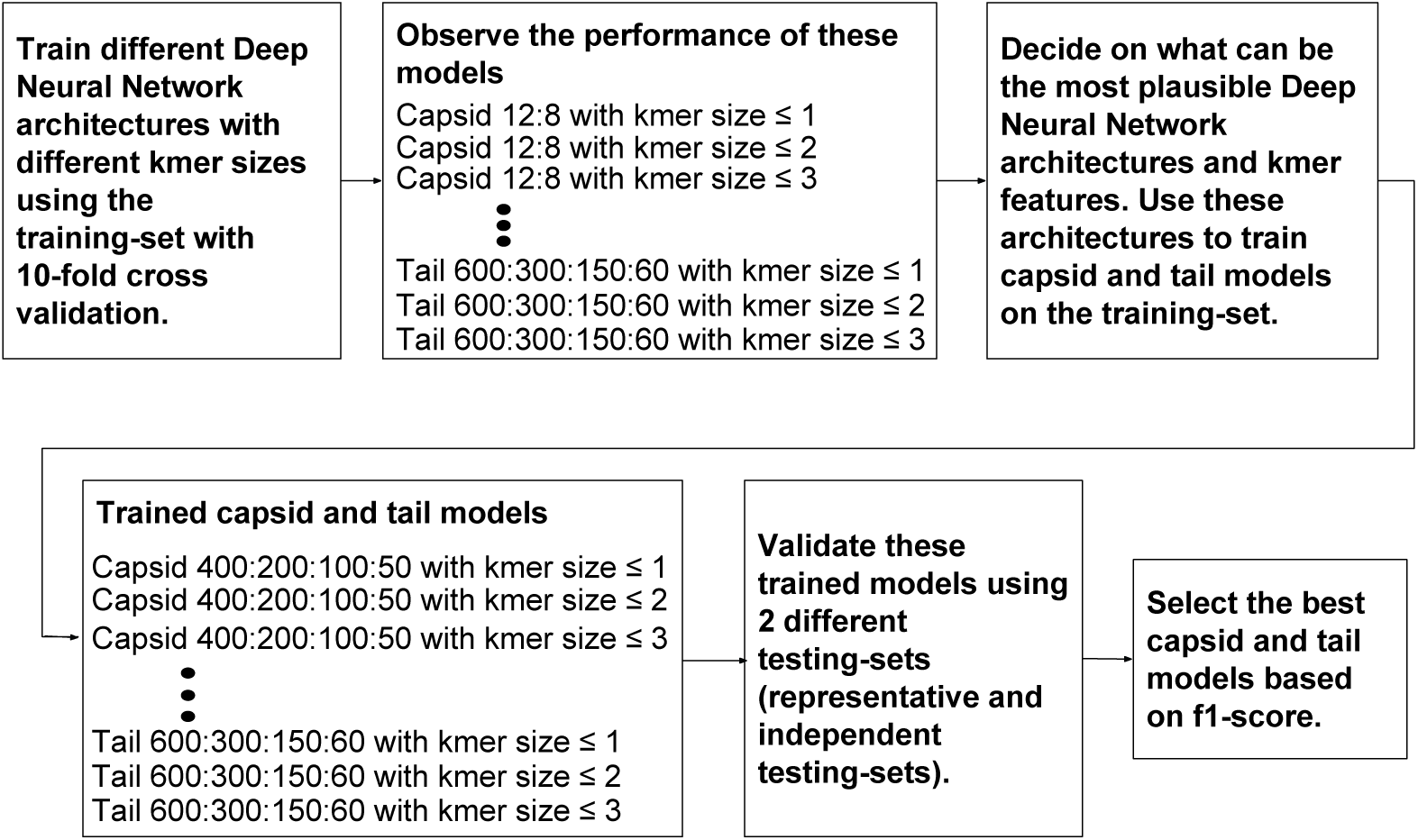
The four steps of DeepCapTail pipeline. (1) decide on the most effective deep neural network architecture and *k*-mer features using the performance of 10-fold cross validation; (2) use these most effective architectures and features to train capsid and tail models using the training set; (3) validate these models using two different testing sets: representative and independent testing sets; (4) select the best capsid and tail models based on the F1-score.

### Performance of Capsid and Tail Models on The Training set

Initially it was unclear what would be an applicable deep learning architecture for this problem. For this reason, different deep learning architectures were examined and compared on the training set with 10-fold cross-validation. First, an extremely small deep learning architecture with two layers of 8 and 12 nodes was examined; then, the number of layers as well as the number of nodes was increased until observing an improvement in capsid and tail predictions with the architecture 400:200:100:50 that has four layers. From this, we kept increasing the number of nodes to have the architectures 600:300:150:60 and 800:400:200:100. We also increased the number of layers to five and six, but the performance dropped and therefore we stopped at the architectures with four layers.

Figures 5a and 5b show the F1-scores of capsid and tail models trained using 10-fold cross-validation on the training set. These models are not exceedingly large, and thus training was done in a timely fashion (e.g., training ‘Capsid 400:200:100:50, ≤ 3’ with 10-fold cross-validation took less than 30 minutes using a high performance computing system: Intel’s Broadwell processors (2 x E5-2683v4 2.1GHz) with 128 GB of memory (2400 MHz) and 32 cores). F1-score was reported instead of the accuracy because F1-score is more reliable when the positive and negative classes are imbalanced, which is the case of our data. The next sections show the performance analysis of all the models presented in Figures 5a and 5b.

**Fig 5.**
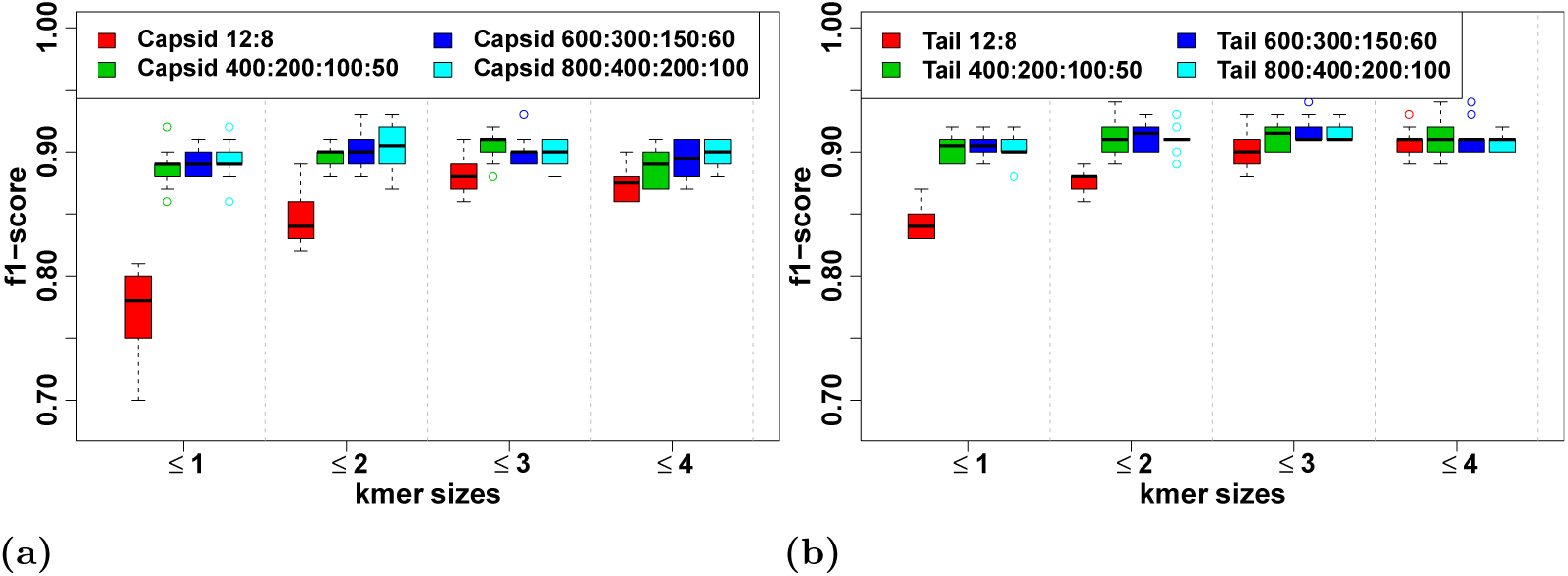
Box-plots of F1-scores of the capsid and tail models using different. *k***-mer features.** *k*-mer features are not mutually exclusive, in other words, a model trained with *k*-mer ≤ 2 has a total of *k*-mer features equal to 420 coming from a total of 20 *k*-mers of size one and 400 *k*-mer of size two. (a) for capsid models and (b) for tail models.

All the models that use *k*-mer size ≤ 4 have either the same or lower F1-scores compared to at least one of the models that use lower *k*-mer sizes and therefore were not considered further for the remaining study (e.g., Figure 5a indicates that the F1-score median of ‘Capsid 400:200:100:50, ≤ 3’ is 92%, whereas, the F1-score median of ‘Capsid 400:200:100:50, ≤ 4’ drops by 3%. For tail models, Figures 5b shows that the F1-score median of ‘Tail:400:200:100:50, ≤ 4’ drops by 1% compared to ‘Tail:400:200:100:50, ≤ 3’). Similarly, the models that have architecture of 12:8 provide the lowest F1-scores compared to all other models and therefore were also not considered in the remaining study (e.g., Figure 5a shows that the F1-score median of ‘Capsid 12:8, ≤ 3’ is 88% compared to 91% for ‘Capsid 400:200:100:50, ≤ 3’. For tail prediction, Figure 5b indicates that ‘Tail 12:8, ≤ 3’ has an F1-score median of 90% compared to 92% for the model ‘Tail 400:200:100:50, ≤ 3’). One can argue that the complexity of the models 400:200:100:50 compared to 12:8 may not justify the little improvement of F1-scores. However, although the models 400:200:100:50 have a higher number of nodes compared to 12:8, they still can be trained in minutes. It takes less than 30 minutes to train ‘Capsid 400:200:100:50, ≤ 3’. The training step is performed once, then the model is saved to predict capsid proteins in just a few milliseconds. Overall, the models 800:400:200:100 have either similar or lower performance than the smaller models 600:300:150:60 and 400:200:100:50 and therefore were not considered further in the remaining study.

To summarize, the models 400:200:100:50 and 600:300:150:60 performed better than 12:8 and 800:400:200:100 as well as the models that use *k*-mer size ≤ 4. For this reason, models with the architectures 400:200:100:50 and 600:300:150:60 using *k*-mer sizes ≤ 1, ≤ 2, and ≤ 3 were further analyzed. In the next section, these models were evaluated using two different testing sets: the representative and independent testing sets.

### Performance of Capsid and Tail models on The Testing sets

In this part of the study, we trained the deep neural network architectures 400:200:100:50 and 600:300:150:60 with different *k*-mer sizes ≤ 1, ≤ 2, and ≤ 3. Then we evaluated these trained models with two different testing sets: the representative and independent testing sets.

#### Using The Representative Testing set

The representative testing set includes sequences that are highly identical to the training set. It assesses the capsid and tail models when the input sequence happens to be similar to the training sequences. Figures 6a and 6b show the ROC curves of capsid and tail models using the representative testing set.

**Fig 6.**
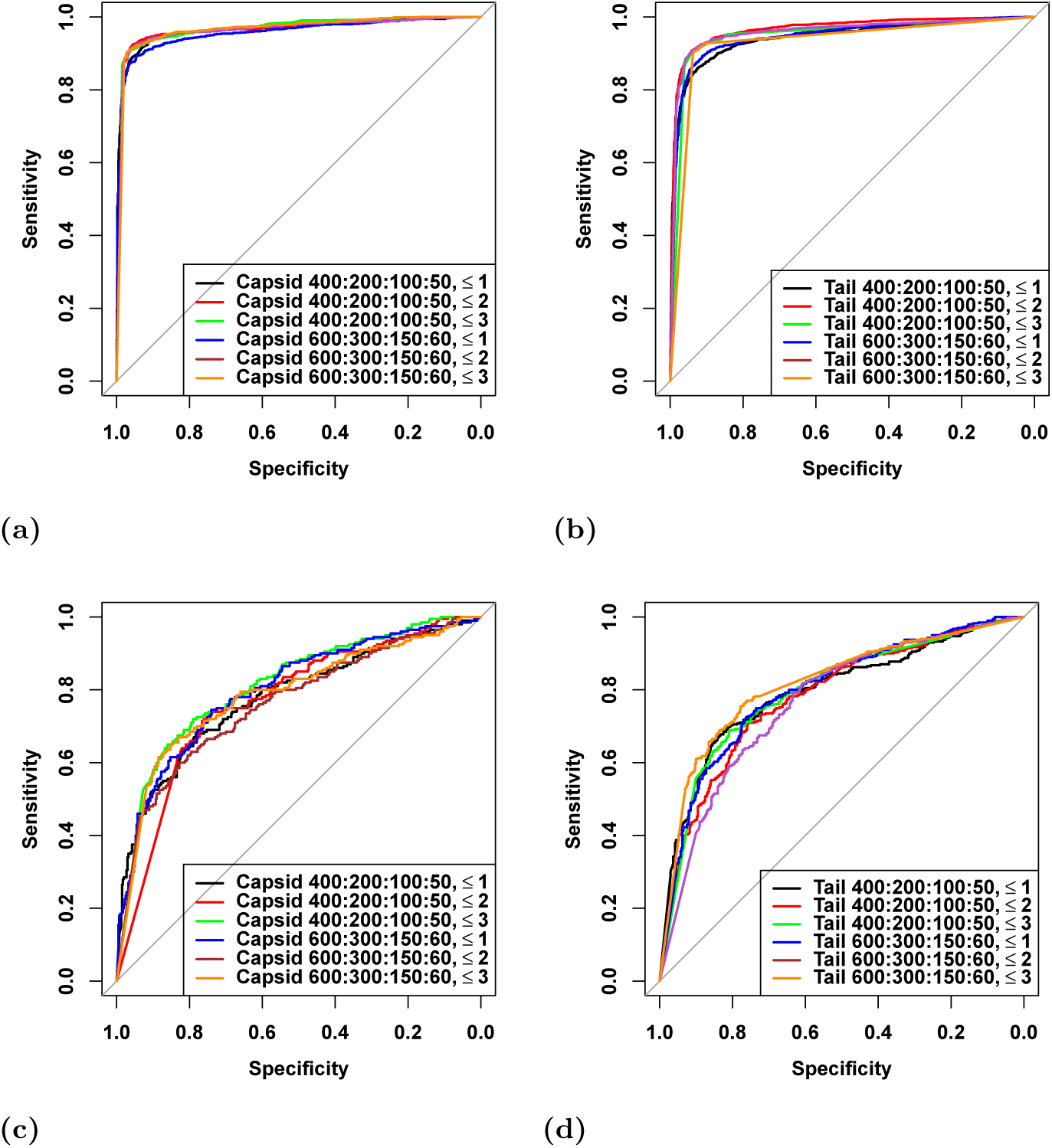
ROC curves of capsid and tail models using the representative and independent testing sets. Figures 6a and 6b are for the evaluation of capsid and tail models using the representative testing set. Figures 6c and 6d are for the evaluation of capsid and tail models using the independent testing set.

Figure 6a shows that the ROC curves of the different capsid models are extremely similar: their AUCs are between 96% and 97%. Figure 6b shows that the ROC curves of the tail models are similar as well: their AUCs are between 93% and 97%. The AUCs of capsid and tail models using the representative testing set are both greater than 90%. A cut-off of 0.5 was used to compute the accuracy, F1-score, recall, and precision of these models; Table 3 shows the results.

**Table 3.**
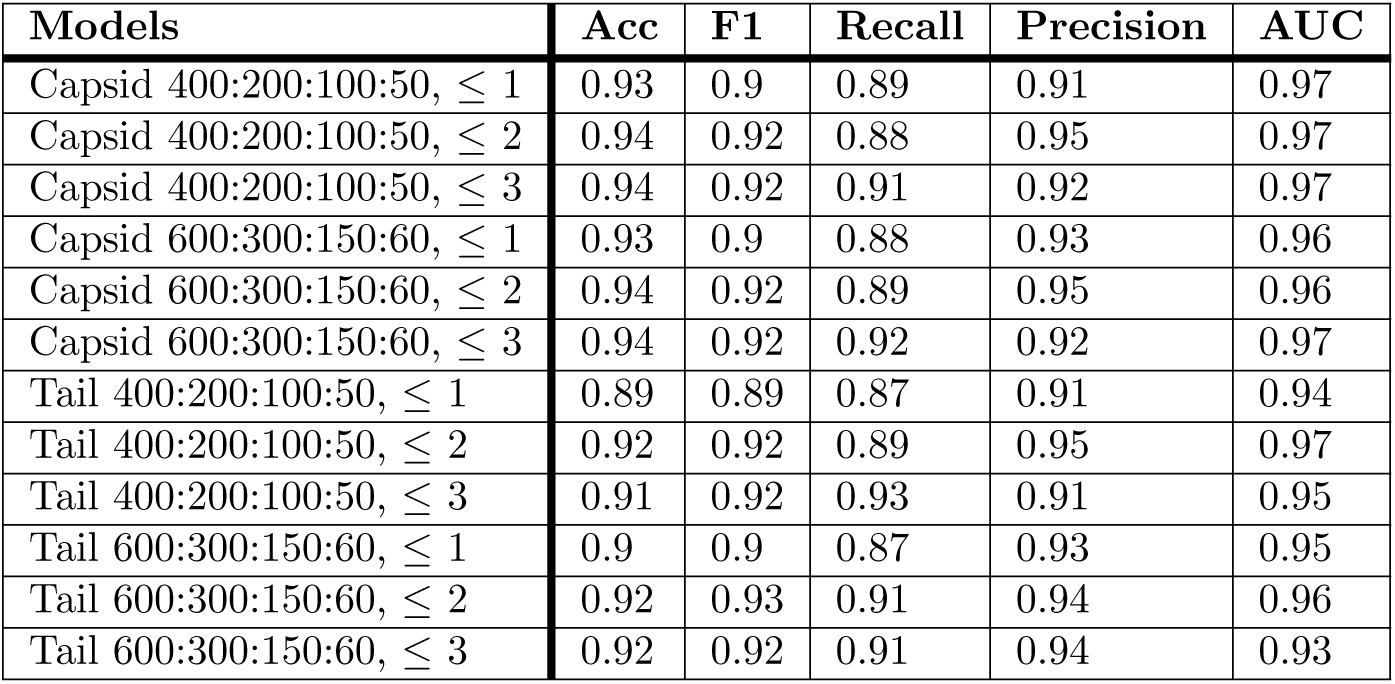
Evaluation of Accuracy, F1-score, Recall, Precision, and AUC of capsid and tail models using the representative testing set.

| Models | Acc | F1 | Recall | Precision | AUC |
| --- | --- | --- | --- | --- | --- |
| Capsid 400:200:100:50, $\leq 1$ | 0.93 | 0.9 | 0.89 | 0.91 | 0.97 |
| Capsid 400:200:100:50, $\leq 2$ | 0.94 | 0.92 | 0.88 | 0.95 | 0.97 |
| Capsid 400:200:100:50, $\leq 3$ | 0.94 | 0.92 | 0.91 | 0.92 | 0.97 |
| Capsid 600:300:150:60, $\leq 1$ | 0.93 | 0.9 | 0.88 | 0.93 | 0.96 |
| Capsid 600:300:150:60, $\leq 2$ | 0.94 | 0.92 | 0.89 | 0.95 | 0.96 |
| Capsid 600:300:150:60, $\leq 3$ | 0.94 | 0.92 | 0.92 | 0.92 | 0.97 |
| Tail 400:200:100:50, $\leq 1$ | 0.89 | 0.89 | 0.87 | 0.91 | 0.94 |
| Tail 400:200:100:50, $\leq 2$ | 0.92 | 0.92 | 0.89 | 0.95 | 0.97 |
| Tail 400:200:100:50, $\leq 3$ | 0.91 | 0.92 | 0.93 | 0.91 | 0.95 |
| Tail 600:300:150:60, $\leq 1$ | 0.9 | 0.9 | 0.87 | 0.93 | 0.95 |
| Tail 600:300:150:60, $\leq 2$ | 0.92 | 0.93 | 0.91 | 0.94 | 0.96 |
| Tail 600:300:150:60, $\leq 3$ | 0.92 | 0.92 | 0.91 | 0.94 | 0.93 |

Table 3 shows that the capsid and tail models performed exceptionally well on the represented testing set (e.g., F1-scores are equal or higher than 89%). The models that use *k*-mer size ≤ 2 or ≤ 3 outperformed the models that use *k*-mer size ≤ 1 (e.g., the F1-score of Capsid 400:200:100:50 using *k*-mer size ≤ 2 and ≤ 3 is 92% compared to 90% for the same architecture using *k*-mer size ≤ 1). However, it is difficult to know if the models that use *k*-mer size ≤ 3 are better than the models that use *k*-mer size ≤ 2 since they have the same F1-score. These observations are valid for both capsid and tail models. The next section shows the details on how these models performed differently on the independent testing set.

#### Using The Independent Testing set

The independent testing set is another testing set used to assess the capsid and tail models. Contrarily to the representative testing set, the independent testing set is substantially different from the training set. It evaluates the capsid and tail models when the input sequence happens to be highly divergent from the training sequences, which might be the case of many newly sequenced phage proteins.

Figures 6c and 6d show the ROC curves of capsid and tail models using the independent testing set. The performance of these models dropped, which is expected because the sequences of the independent testing set are highly divergent from the sequences of the training set (e.g., the AUC of ‘Capsid 400:200:100:50, ≤ 3’ dropped from 97% to 81%. The AUC of ‘Tail 600:300:150:60, ≤ 3’ dropped from 93% to 82%). Both capsid and tail AUCs are above 80%.

We compute the accuracy, F1-score, recall, and precision of these models using a cut-off of 0.5. Table 4 shows the results.

**Table 4.** Evaluation of Accuracy, F1-score, Recall, Precision, AUC of capsid and tail models using the independent set.

| Models | Acc | F1 | Recall | Precision | AUC |
| --- | --- | --- | --- | --- | --- |
| Capsid 400:200:100:50, $\leq 1$ | 0.77 | 0.61 | 0.52 | 0.71 | 0.78 |
| Capsid 400:200:100:50, $\leq 2$ | 0.69 | 0.62 | 0.75 | 0.53 | 0.77 |
| Capsid 400:200:100:50, $\leq 3$ | **0.79** | **0.67** | **0.64** | **0.7** | **0.81** |
| Capsid 600:300:150:60, $\leq 1$ | 0.78 | 0.6 | 0.5 | 0.75 | 0.8 |
| Capsid 600:300:150:60, $\leq 2$ | 0.76 | 0.6 | 0.52 | 0.69 | 0.76 |
| Capsid 600:300:150:60, $\leq 3$ | 0.73 | 0.64 | 0.72 | 0.57 | 0.79 |
| Tail 400:200:100:50, $\leq 1$ | 0.75 | 0.72 | 0.66 | 0.81 | 0.79 |
| Tail 400:200:100:50, $\leq 2$ | 0.7 | 0.73 | 0.81 | 0.66 | 0.78 |
| Tail 400:200:100:50, $\leq 3$ | 0.68 | 0.73 | 0.88 | 0.62 | 0.79 |
| Tail 600:300:150:60, $\leq 1$ | 0.74 | 0.74 | 0.72 | 0.75 | 0.8 |
| Tail 600:300:150:60, $\leq 2$ | 0.7 | 0.71 | 0.75 | 0.68 | 0.77 |
| Tail 600:300:150:60, $\leq 3$ | **0.76** | **0.76** | **0.76** | **0.77** | **0.82** |

Best capsid and tail models are in bold.

Contrary to the results of the representative testing set, it is easier to distinguish the best models for capsid and tail predictions using the independent testing set. The best capsid model is ‘Capsid 400:200:100:50, ≤ 3’, with an accuracy and F1-score of 76% and 66% respectively. The best tail model is ‘Tail 600:300:150:60, ≤ 3’, with an accuracy and F1-score of 76%. We compare our best models with state-of-the-art prediction programs in the next section.

### Performance Comparison of Our Best Capsid and Tail Models with State-Of-The-Art using Two Different Testing sets

We compare our best capsid model ‘Capsid 400:200:100:50, ≤ 3’ and our best tail model ‘Tail 600:300:150:60, ≤ 3’ with two state-of-the-art programs, iVIREONS [7] and VIRALpro [8]. To this end, the representative and independent testing sets were used and results are shown in Figure 7.

**Fig 7.**
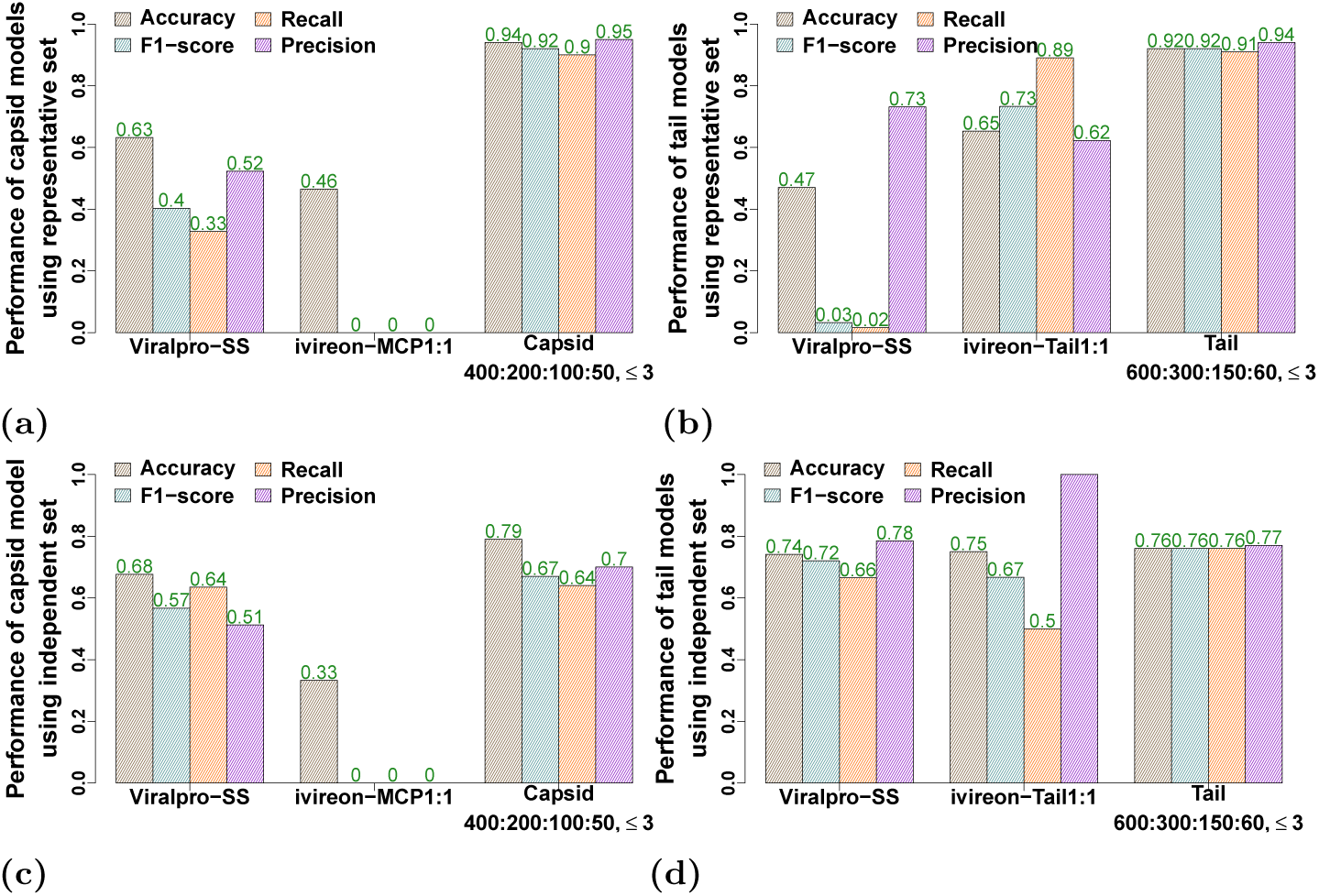
Barplot of the accuracy, F1score, recall, and precision of our best capsid and tail models as well as iVIREONS and VIRALpro. Figures 7a and 7b evaluate the capsid and tail models using the representative testing set. Figures 7c and 7d evaluate the capsid and tail models using the independent testing.

Our capsid model ‘Capsid 400:200:100:50, ≤ 3’ and our tail model ‘Tail 600:300:150:60, ≤ 3’ outperform iVIREONS [7] and VIRALpro [8] with both the representative and independent testing sets. Using the representative testing set, our capsid model ‘Capsid 400:200:100:50, ≤ 3’ has a F1-score of 92% compared to 0% and 40%, and our tail model ‘Tail 600:300:150:60, ≤ 3’ records a F1-score of 92% compared to 73% and 0.03% for iVIREONS and VIRALpro respectively.

The successful prediction of most of the sequences in the representative testing set is expected as our models were trained on similar sequences. However, iVIREONS and VIRALpro were trained on sequences that are different from the representative testing set. Supplemental Figure 2 shows that more than 60% of the training set of the two models have an identity less than 40% to the representative testing set, which can be the reason why they were unable to perform as well as our models.

However, using the independent testing set, the performance of our models dropped, but still performed better than iVIREONS and VIRALpro. Our capsid model ‘Capsid 400:200:100:50, ≤ 3’ outperforms iVIREONS and VIRALpro with an F1-score of 67% compared to 0% and 57% respectively, and our tail model ‘Tail 600:300:150:60, ≤ 3’ presents an F1-score equal to 76% compared to 67% and 72% for iVIREONS and VIRALpro respectively.

The performance of our models dropped with the independent testing set because the testing set is substantially different from the sequences used to train the models. The independent testing set is also different from the training set used by both iVIREONS and VIRALpro.

Supplemental Figure 3 shows that more than 70% of the training set used by these models have an identity less than 40% to this testing set. This means all of these models are tested on sequences that are different from their training sequences, and for this reason, we consider the independent testing set unbiased compared to the representative testing set.

### Performance Comparison of Our Best Capsid and Tail Models with State-Of-The-Art using Viral Metagenomic Data

We compare the performance of ‘Capsid 400:200:100:50, ≤ 3’ and ‘Tail 600:300:150:60, ≤ 3’ with iVIREONS [7] and VIRALpro [8] using viral metagenomic data. We use the same viral metagenomic data that were employed by VIRALpro [8] to assess their capsid and tail predictors. This data consists of five different metagnomic datasets with no homology to known proteins. These five datasets do not have any capsid or tail annotation, and this is why we cannot compute F1-scores on this dataset. We present relevant details about these datasets in Table 5 (more details on these datasets can be found in [8]).

**Table 5.** Overview of the 5 metagenomic datasets that we use to compare our capsid and tail models with the iVIREONS and VIRALpro.

| Metagenomic datasets | # of sequences | Relevant details |
| --- | --- | --- |
| Moore phages | 1172 | marine phage proteins. |
| RnaCoastal | 86 | RNA viral proteins of sea water samples. |
| Oresund-Struct | 85 | sea water phages isolated from the Strait of Oresund; using MS-based proteomics this dataset are identified as viral structural proteins. |
| Oresund-Hypo | 524 | sea water phages isolated from the Strait of Oresund; using MS-based proteomics this dataset are identified as viral hypothetical proteins. |
| Oresund-NonStruct | 156 | sea water phages isolated from the Strait of Oresund; using MS-based proteomics this dataset are identified as viral non-structural proteins. |

We present in Figure 8 the Venn Diagram of capsid and tail predictions using the metagenomic dataset of Oresund Struct. As we detailed in Table 5, this dataset is identified as structural viral proteins by MS-based proteomics [8]. For tail prediction, our model as well as iVIREONS and VIRALpro agreed on the prediction of most of the tail sequences: 33 sequences were identified by all these models as tail sequences.

**Fig 8.**
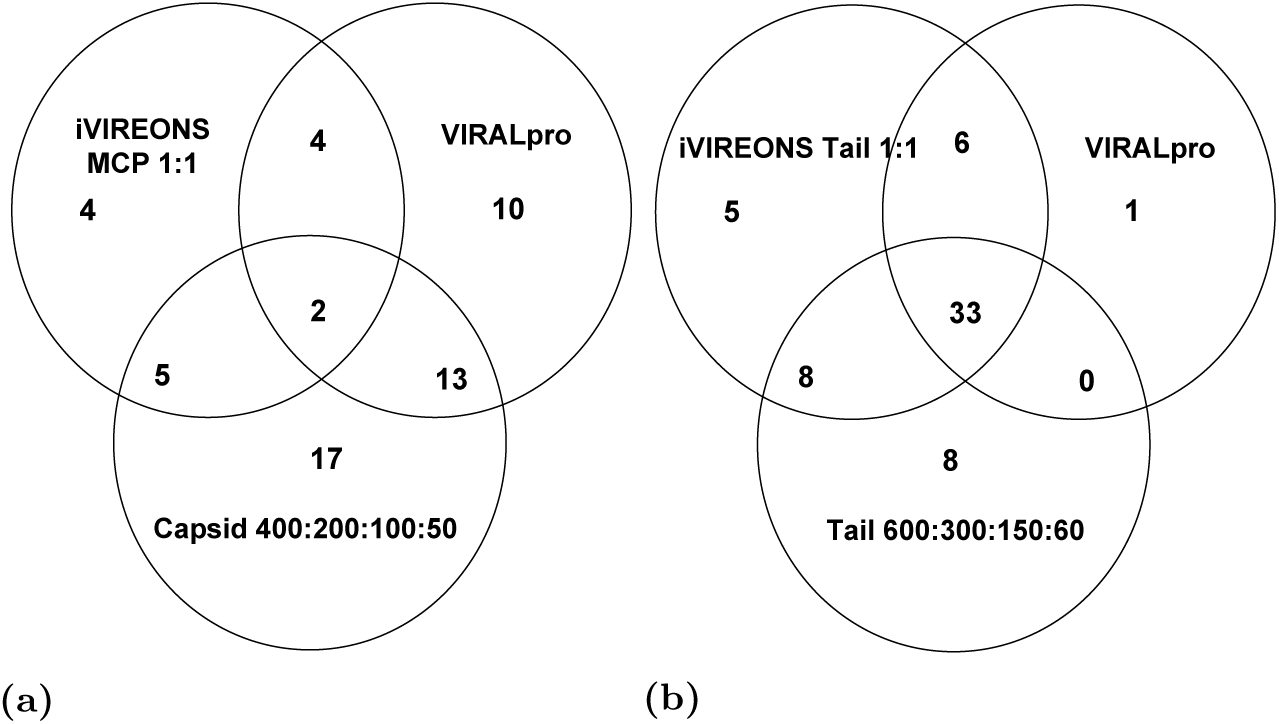
Venn Diagram of the capsid and tail predictions of our models as well as iVIREONS and VIRALpro using the metagenomic dataset ‘Oresund Struct’. Figure 8a presents the number of predicted capsid proteins, and Figure 8b presents the number of predicted tail proteins

However, for the capsid prediction, our model and the iVIREONS and VIRALpro agreed on the prediction of only two capsid sequences. It is difficult to know if these predictions are correct, since we do not have the annotation of capsid and tail proteins for this metagenomic dataset. The Venm Diagram of the prediction of capsid and tail proteins for the four remaining of metagenomic datasets is shown in the supplemental material.

## Conclusion

We proposed the deep learning models ‘Capsid 400:200:100:50, ≤ 3’ and ‘Tail 600:300:150:60, ≤ 3’ that predicts capsid and tail proteins of phages. We evaluated these models using two different testing sets. Our models outperformed the state-of-the-art iVIREONS and VIRALpro, which suggests that our models are more accurate in prediction. We also compared the performance of our models, iVIREONS and VIRALpro using five different viral metagenomic datasets. All of these models agreed on the annotation of some of the capsid and tail proteins; however, it is difficult to assess the accuracy of these models since the correct answer is not known.

